# The Impact of Cross-Species Gene Flow on Species Tree Estimation

**DOI:** 10.1101/820019

**Authors:** Xiyun Jiao, Thomas Flouris, Bruce Rannala, Ziheng Yang

**Author notes:** Ziheng Yang, Department of Genetics, University College London, Gower Street, London WC1E 6BT, UK.

## Abstract

Recent analyses of genomic sequence data suggest cross-species gene flow is common in both plants and animals, posing challenges to species tree inference. We examine the levels of gene flow needed to mislead species tree estimation with three species and either episodic introgressive hybridization or continuous migration between an outgroup and one ingroup species. Several species tree estimation methods are examined, including the majority-vote method based on the most common gene tree topology (with either the true or reconstructed gene trees used), the UPGMA method based on the average sequence distances (or average coalescent times) between species, and the full-likelihood method based on multi-locus sequence data. Our results suggest that the majority-vote method is more robust to gene flow than the UPGMA method and both are more robust than likelihood assuming a multispecies coalescent (MSC) model with no cross-species gene flow. A small amount of introgression or migration can mislead species tree methods if the species diverged through speciation events separated by short time intervals. Estimates of parameters under the MSC with gene flow suggest the *Anopheles gambia* African mosquito species complex is an example where gene flow greatly impacts species phylogeny.

---

Cross-species hybridization or introgression has long been recognized as a process that can generate biological diversity, especially in plants (e.g., Anderson, 1949; Mallet, 2007). Analyses of genomic data in the past few years suggest that introgression is also common in animals (Mao *et al*., 2018; Ellegren *et al*., 2012; Kumar *et al*., 2017; Wu *et al*., 2018; Chan *et al*., 2013), including humans and their close relatives (Nielsen *et al*., 2017). Introgression may involve either sister or non-sister species (e.g., Mallet *et al*., 2016) and may play an important role in the speciation process (Mallet *et al*., 2016; Martin and Jiggins, 2017). Introgression, together with deep coalescence (also known as incomplete lineage sorting, ILS), may cause difficulties for species tree reconstruction. In extreme cases, the whole genome, and in particular the autosomes, are affected by such pervasive gene flow that they do not reflect the species phylogeny. This appears, for example, to be the case with the *Anopheles gambiae* species complex, in which the autosomes suggest different species relationships from the X chromosome, with the X chromosome reflecting the true history of species divergences (Fontaine *et al*., 2015; Thawornwattana *et al*., 2018). The *Heliconius* butterflies appear to be another such case, with the Z chromosome favoring different phylogenies from the autosomes (Edelman *et al*., 2018). In both examples, the species arose through a rapid succession of speciation events, generating very short interior branches in the phylogeny.

A number of methods have been developed to detect gene flow across species using genetic sequence data, including population genetic methods based on *F*_st_ and summary methods that make use of observed site patterns (Green *et al*., 2010; Durand *et al*., 2011) or estimated gene tree topologies (Yu and Nakhleh, 2015). Full-likelihood methods based on sequence alignments (Hey and Nielsen, 2004; Hey *et al*., 2018; Wen and Nakhleh, 2018; Zhang *et al*., 2018) are also being actively developed. See Degnan (2018) and Folk *et al*. (2018) for recent reviews.

Here we consider the question of how much gene flow is sufficient to mislead species tree estimation methods that accommodate the coalescent process but not gene flow. As discussed by Folk *et al*. (2018), the impact of gene flow on the tree of life is an important topic worth serious study. We focus on the case of three species with sequences evolving under the molecular clock and are specifically interested in closely related species, for which gene flow may be a major concern. We consider two distinct models of gene flow, both of which accommodate the multispecies coalescent (MSC). The isolation-with-migration (IM) model assumes continuous migration, with the species exchanging migrants at certain rates every generation (Hey and Nielsen, 2004; Hey, 2010). The multispecies-coalescent-with-introgression (MSci) model assumes episodic introgression/hybridization events (Degnan, 2018; Yu *et al*., 2014). We consider two scenarios of gene flow on the species tree (*A*, (*B,C*)). Inflow is introgression (or migration) from the outgroup species *A* to the ingroup species *C*, and outflow is in the reverse direction. In both scenarios, gene flow makes species *A* and *C* look similar, potentially misleading species tree methods to infer the incorrect tree (*B*, (*C, A*)).

Previously Leaché *et al*. (2014) used computer simulation to examine the impact of continuous migration on the estimation of the species tree, finding that the effects depend on the species involved in gene flow. Solis-Lemus *et al*. (2016) used simulation to study the statistical consistency of several species tree methods including ASTRAL (Mirarab *et al*., 2014) and NJST (Liu and Yu, 2011), which treat unrooted gene tree topologies as data. A method was considered inconsistent if the probability of retrieving the correct species tree fails to increase when the number of gene trees (the number of loci) increases. The approach we take here is largely analytical, not affected by sampling errors in simulation. Hahn and Nakhleh (2016, fig. 3) calculated gene-tree probabilities under an introgression model, arguing that the concept of a species tree is poorly defined when there is gene flow. Long and Kubatko (2018) studied the probabilities of gene tree topologies under an isolation-with-initial-migration model (Wilkinson-Herbots, 2012) for three species, in which there is initial gene flow between sister species after their divergence. One might expect gene flow between sister species to make them appear more similar, making it easier to infer the species tree, but surprisingly with different population sizes, gene flow between sister species can cause the most probable gene tree topology to differ from the species tree, leading to so-called anomalous gene trees. Here we consider gene flow affecting non-sister species and both continuous migration and episodic introgression. We consider several species tree inference methods, including the majority-vote method based on the true gene trees studied by Long and Kubatko (2018).

**Fig. 1.**
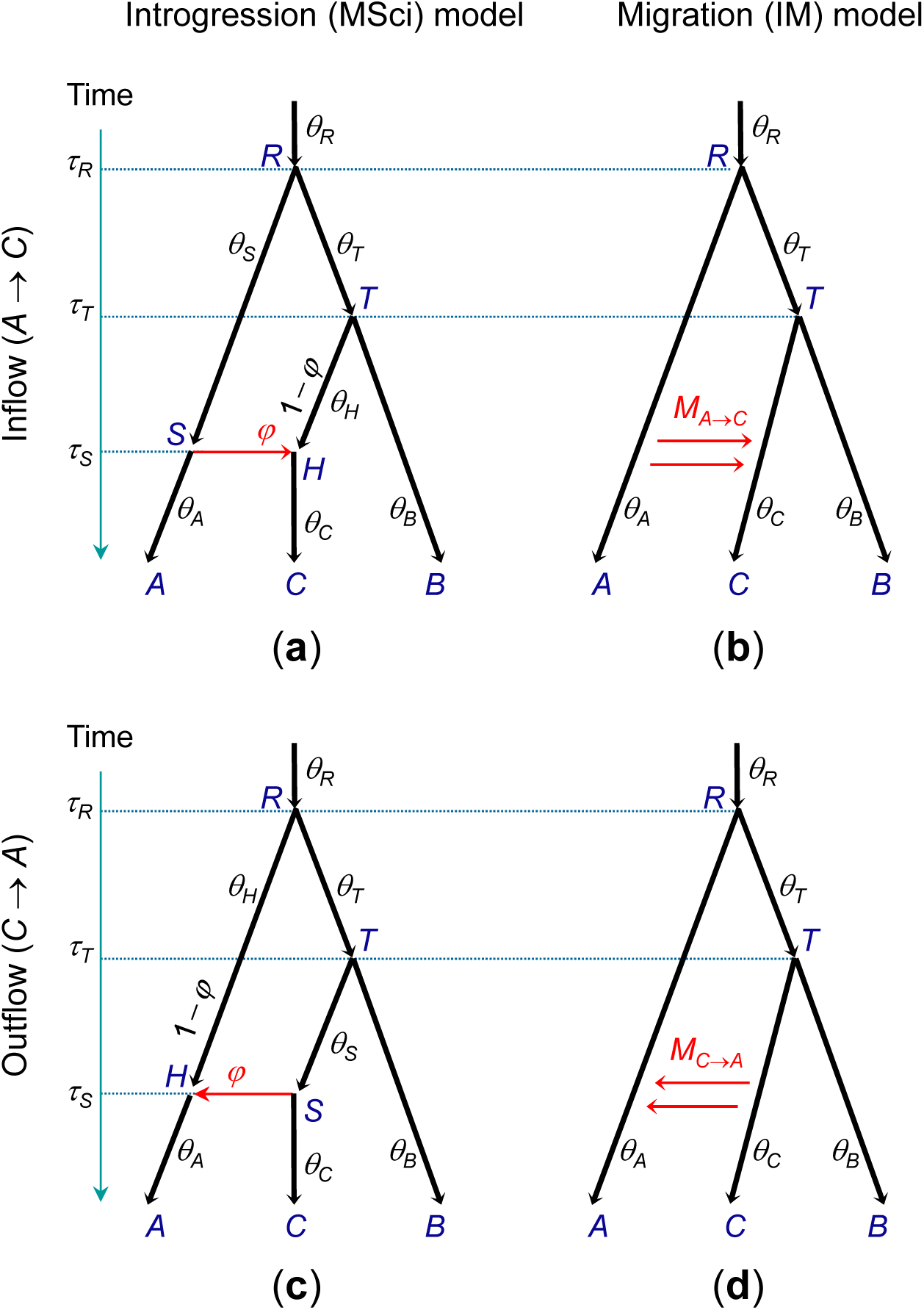
Species tree (*A*, (*B,C*)), with introgression (**a** and **c** or migration (**b** and **d**) from the outgroup species *A* to the ingroup species *C* (inflow, **a** and **b**) or in the reverse direction (outflow, **c** and **d**).

**Fig. 2.**
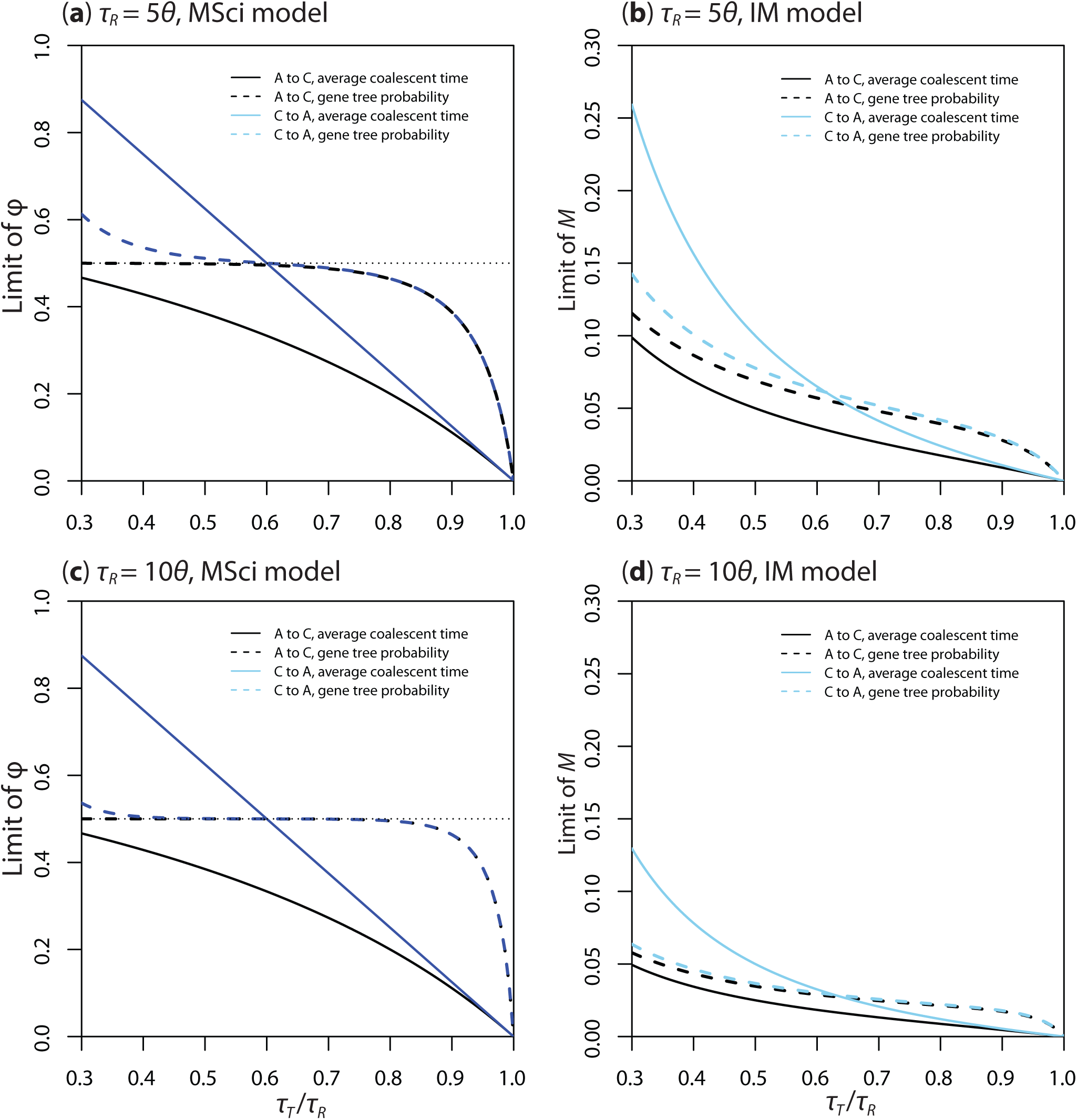
The lower limit of the introgression probability *φ* in the MSci model and of the migration rate *M* in the IM model that is necessary and sufficient to mislead species tree estimation, plotted against *τ*_*T*_ /*τ*_*R*_. All populations are assumed to have the same size parameter (*θ*), while the age of the root is *τ*_*R*_ = 5*θ* in (a) and (b) and *τ*_*R*_ = 10*θ* in (c) and (d). In the MSci model, we use *τ*_*H*_ = *τ*_*S*_ = *τ*_*R*_/5. Note that the *φ* and *M* limits do not depend on the precise value of *θ*.

**Fig. 3.**
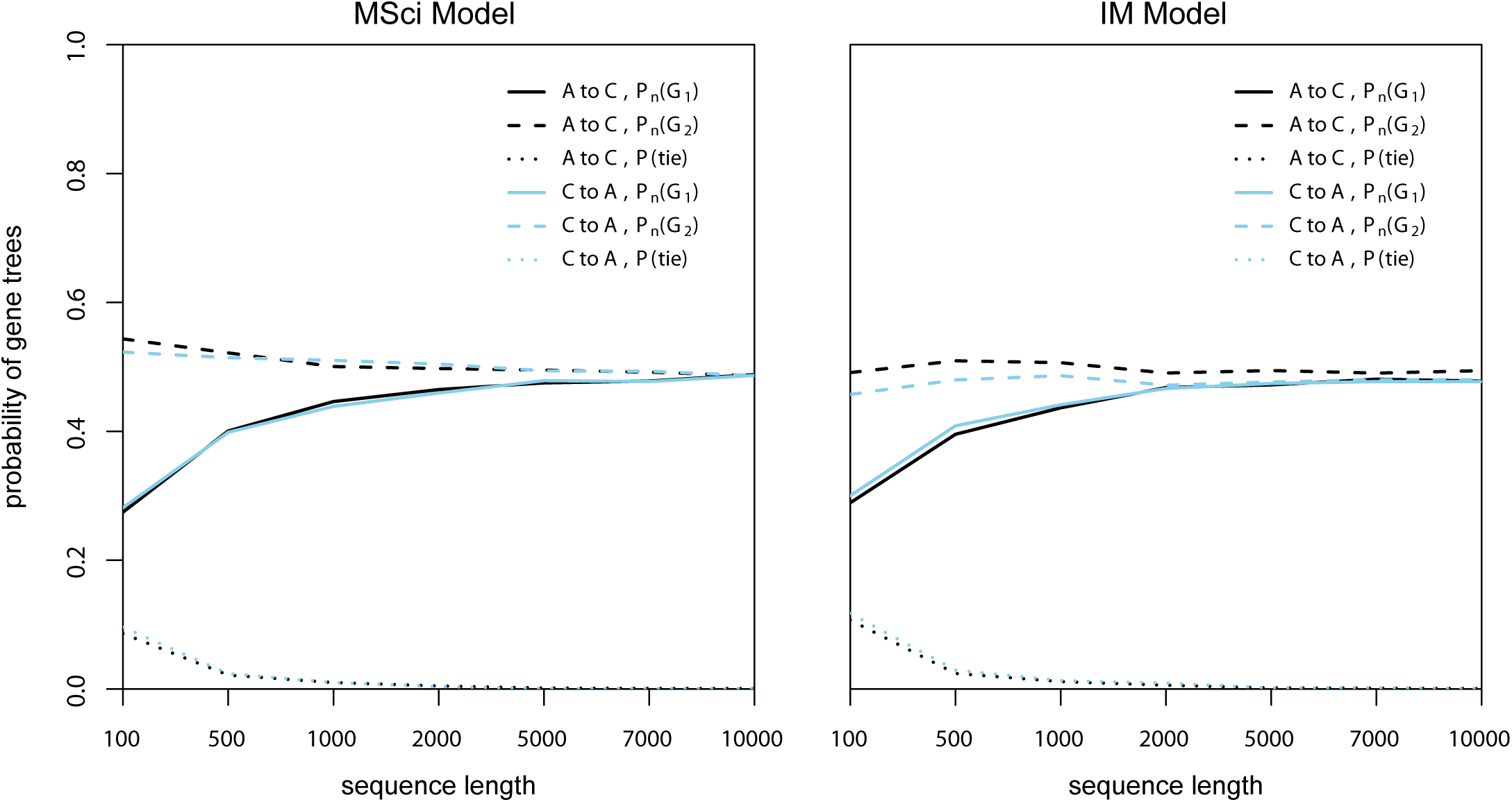
Probabilities of estimated gene trees, ℙ_*n*_(*G*_1_) and ℙ_*n*_(*G*_2_), as a function of sequence length (*n*), under the MSci and IM models. The following parameter values are used: *θ* = 0.01 for all populations, *τ*_*T*_ = 0.04, and *τ*_*R*_ = 0.05; and for the MSci model, *τ*_*H*_ = 0.01 and *φ* = 0.4638 for inflow introgression (*A* → *C*) and 0.4643 for outflow introgression (*C* → *A*), while for the IM model, *M* = 0.0393 for inflow migration and 0.0419 for outflow immigration. At those parameter values, ℙ(*G*_1_) = ℙ(*G*_2_) when the sequence length is infinity (so that there are no phylogenetic reconstruction errors). ℙ(tie) is the proportion of datasets in which two or three gene trees are equally best.

The species tree estimation methods considered here are all statistically consistent in the case of three species when there is deep coalescence but no gene flow. First, we study the majority-vote method of using the most common (true) gene tree topology as the estimate of the species tree. This is known to be statistically consistent in the case of three species and three sequences per locus when there is no gene flow, with the estimated species tree approaching the true tree when the number of loci approaches infinity (Hudson, 1983). We derive the distribution of the *true* gene tree topologies under each model of gene flow, and examine the impact of phylogenetic reconstruction errors when the *estimated* gene trees are used to estimate the species tree. Use of estimated gene trees is known to be consistent in the case of three species when the model involves coalescent but no gene flow (Yang, 2002). Next we consider the UPGMA method, which uses the average sequence distance between species (or the average coalescent time between species since we assume the molecular clock) to infer the species tree (Liu *et al*., 2009). This is equivalent to calculating sequence distances using concatenated data followed by UPGMA reconstruction of the species tree. This method is known to be consistent in the case of three species (Liu *et al*., 2009). Finally, we consider the ML method of species tree estimation, which averages over the gene trees and branch lengths and thus accounts for phylogenetic reconstruction errors (Xu and Yang, 2016; Yang, 2002). While full-likelihood methods of species tree estimation under the MSC applied to multilocus sequence alignments, including maximum likelihood (ML) (Yang, 2002; Zhu and Yang, 2012) and Bayesian inference (Liu and Pearl, 2007; Heled and Drummond, 2010; Yang and Rannala, 2014), are analytically intractable (Xu and Yang, 2016), the 3S program has an efficient ML implementation that can handle thousands of loci. We thus use 3S to analyze tens of thousands of loci to approximate the case of infinite data. Note that our interest is in the consistency or inconsistency of each species tree estimation method in the face of gene flow as the number of loci approaches infinity. To our knowledge, this represents the first analysis of full-likelihood methods based on sequence alignments, which may be expected to make the most efficient use of information in the sequence data and to have statistically optimal performance when the model is correct. The performance of likelihood methods when the model is mis-specified is unknown.

## 1. THEORY

We consider two gene flow scenarios: “inflow” where there is gene flow (introgression or migration) from the outgroup species *A* to the ingroup species *C* on the species tree (*A*, (*B,C*)) (Fig. 1a & b) and “outflow” in the opposite direction (Fig. 1c & d). Both the species divergence times (*τ*_*R*_, *τ*_*S*_, and *τ*_*T*_) and the population size parameters (*θ* s) in the model are measured by the expected number of mutations/substitutions per site. The data consist of multiple loci, with three sequences – one from each of the three species at each locus (*a, b, c*). The possible gene trees at each locus are *G*_1_ = (*a*, (*b, c*)), which matches the species tree, *G*_2_ = (*b*, (*c, a*)), which reflects the introgression or migration pattern, and *G*_3_ = (*c*, (*a, b*)) which matches neither. Sequence data are then analyzed to infer the species tree under the multispecies coalescent (MSC) model (assuming no gene flow). We derive analytical results for two simple methods of species tree estimation assuming no gene flow: (1) the majority-vote method for which the correct species tree is inferred if ℙ(*G*_1_) > ℙ(*G*_2_) and (2) the UPGMA method using the average coalescent times across loci (Liu *et al*., 2009) for which the correct species tree is inferred if 𝔼(*t*_*bc*_) < 𝔼(*t*_*ac*_). Initially, we ignore sampling errors in the gene trees or the estimated sequence distances. We study the asymptotic behavior of the two methods as the number of loci approaches infinity. We derive results under the two methods for two distinct models of gene flow: instantaneous admixture under the MSci model and ongoing gene flow under the IM model.

### Gene flow under the MSci model

Here, we consider properties of the majority-vote and UPGMA methods for data arising under a MSci model of admixture. Referring to Figure 1 we define

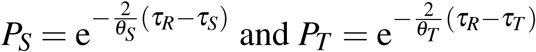

to be the probability that two sequences entering either species *S*, or species *T*, do not coalesce in that species and instead both enter the ancestral species (species *R* for both). Note that 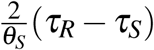 and 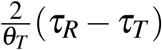 are known as the branch lengths in coalescent units in Fig. 1. We first consider properties of the methods with data generated by inflow.

#### UPGMA Method with Inflow

Here we derive results for a model assuming instantaneous introgression from *A* to *C* (inflow), with the introgression probability *φ*. Sequences *a* and *b* can coalesce in *R* only, and the coalescent time has the exponential density 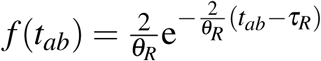 with expectation 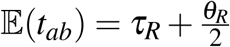. Sequences *a* and *c* can coalesce in population *S* as well as *R*, while sequences *b* and *c* can coalesce in species *T* as well as *R*. This means that 𝔼(*t*_*ab*_) > max{𝔼(*t*_*ac*_), 𝔼(*t*_*bc*_)}. The probability density of the coalescent time *t*_*ac*_ is

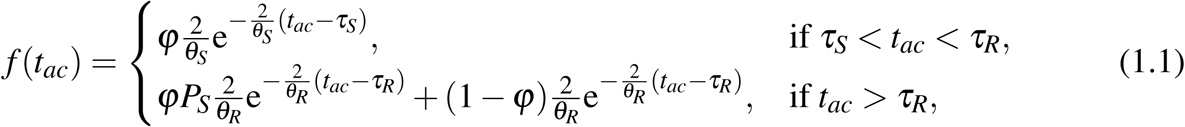

with expectation

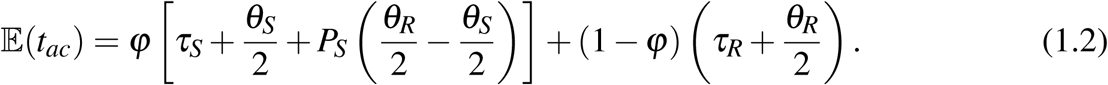

Similarly, sequences *b* and *c* can coalesce in species *T* and *R*, so we have the density

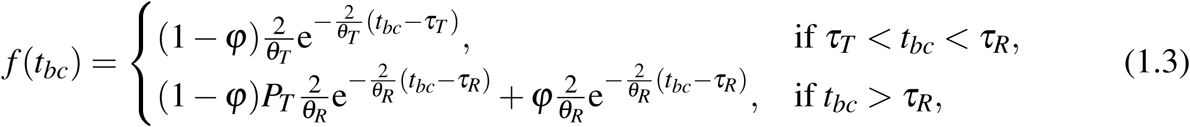

with expectation

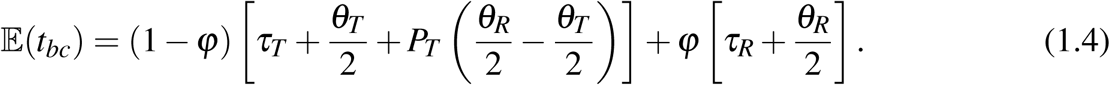

Then 𝔼(*t*_*bc*_) > 𝔼(*t*_*ac*_) and the UPGMA method based on average coalescent times infers an incorrect species tree if and only if

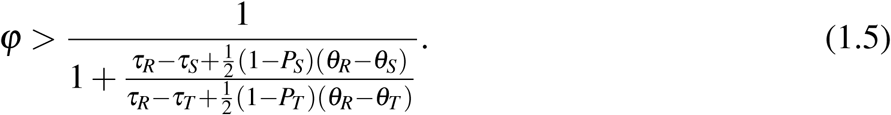

#### Majority-Vote Method with Inflow

If sequences *b* and *c* coalesce in population *T*, the gene tree will be *G*_1_, while if *a* and *c* coalesce in population *S*, the gene tree will be *G*_2_. If neither of those events occurs, both coalescent events for the three sequences will occur in species *R* and the three gene trees will occur with equal probabilities. Thus ℙ(*G*_3_) < min ℙ(*G*_1_), ℙ(*G*_2_). We have

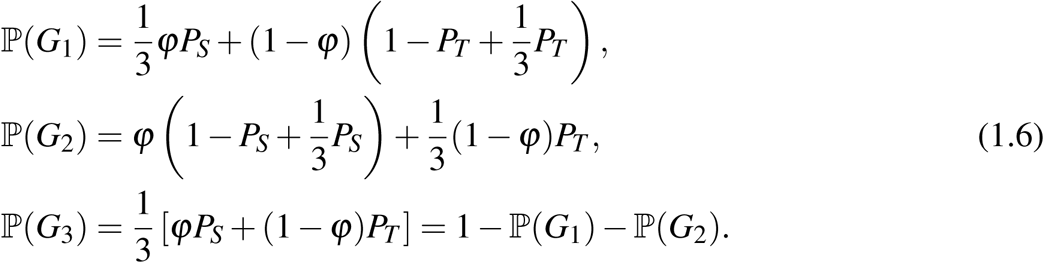

For example, gene tree *G*_1_ results from sequences *b* and *c* coalescing first. If sequence *c* enters species *S* (which happens with probability *φ*), this can occur only if sequences *c* and *a* do not coalesce in species *S*. This is the first term, 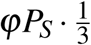, in ℙ(*G*_1_). If sequence *c* enters species *H* (which happens with probability 1 −*φ*),, sequences *b* and *c* can coalesce in species *T* or *R*. Hence the second term 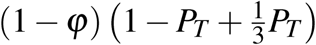.

Thus ℙ(*G*_1_) < ℙ(*G*_2_) and the majority-vote method based on the most common gene tree infers an incorrect species tree if and only if

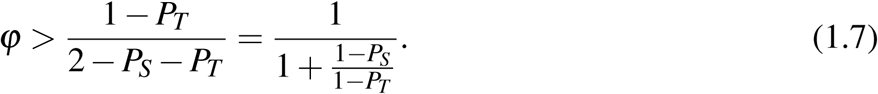

This can also be obtained by noting that ℙ(*G*_1_) < ℙ(*G*_2_) if and only if the probability that sequences *b* and *c* coalesce in population *T* is smaller than the probability that sequences *a* and *c* coalesce in population *S*: that is, if (1 − *φ*)(1 − *P*_*T*_) < *φ*(1 − *P*_*S*_).

Note that the gene tree probabilities (Equation 1.6) and the *φ* limit based on them (Equation 1.7) depend on only the internal branch lengths (in coalescent units) on the species tree, but not the species divergence times and population sizes. The *φ* limits for both gene tree probabilities and the average coalescent times (Equations 1.7 and 1.5) depend on the gene tree topologies and coalescent times but not the mutation rate: are functions of *τ/θ* ratios and not of *τ*s and *θ* s individually.

#### UPGMA Method with Outflow

Here we derive results for a model assuming instantaneous introgression from *C* to *A* (outflow) with introgression probability *φ*. Referring to Figure 1c we define

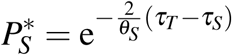

to be the probability that two sequences entering population *S* do not coalesce in that population and instead enter its ancestor (population *T*) (Fig. 1c). Sequences *a* and *c* can coalesce in populations *S, T* and *R*, and sequences *b* and *c* or *a* and *b* can coalesce in species *T* and *R*. The probability density of the coalescent time *t*_*ac*_ is

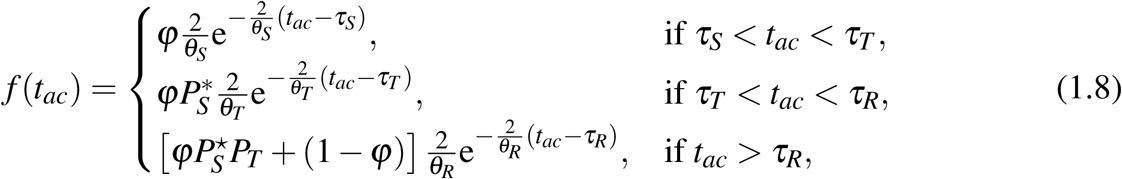

with expectation

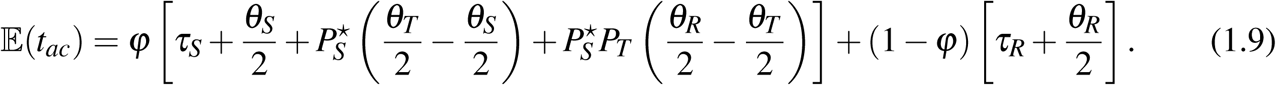

The coalescent time between sequences *b* and *c* has the density

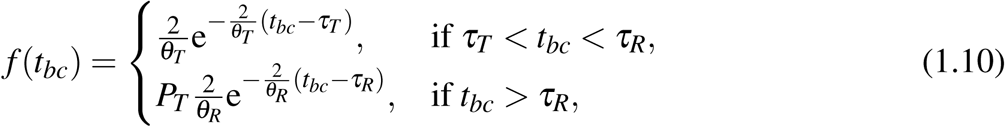

with expectation

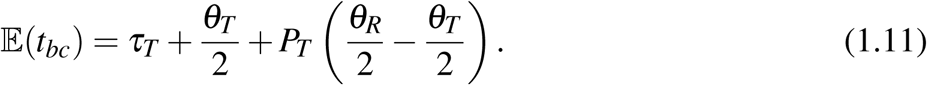

The probability density of the coalescent time *t*_*ab*_ is

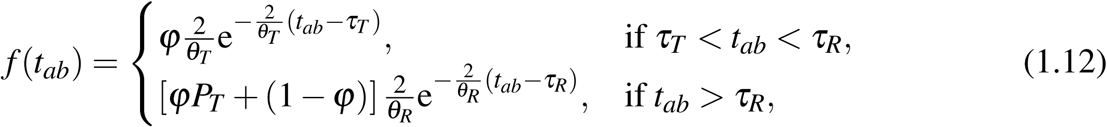

with the expectation

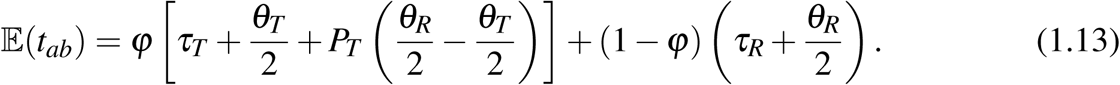

It is easy to see that 𝔼(*t*_*ab*_) > max {𝔼(*t*_*ac*_), 𝔼(*t*_*bc*_)}. Then 𝔼(*t*_*bc*_) > 𝔼(*t*_*ac*_) and the UPGMA method infers an incorrect species tree if and only if

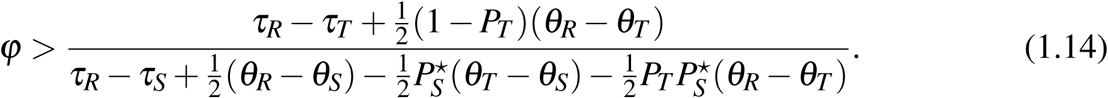

#### Majority-Vote Method with Outflow

When we trace the genealogy of sequences *a, b*, and *c* backward in time, sequence *a* may enter species *S* (with probability *φ*) or species *H* (with probability 1 − *φ*). Consider the first case, of sequence *a* entering species *S*. If *a* and *c* coalesce in population *S*, the gene tree will be *G*_2_. Otherwise, the two coalescent events for the three sequences can occur in either *T* or *R* and the three gene trees occur with equal probabilities. In the second case, sequence *a* enters species *H*. Then if *b* and *c* coalesce in population *T*, the gene tree will be *G*_1_, and otherwise, both coalescent events for the three sequences will occur in *R* with equal probabilities for the three gene trees. This means that *G*_3_ is the least frequent gene tree, with ℙ (*G*_3_) < min {ℙ(*G*_1_), ℙ(*G*_2_)}. We have

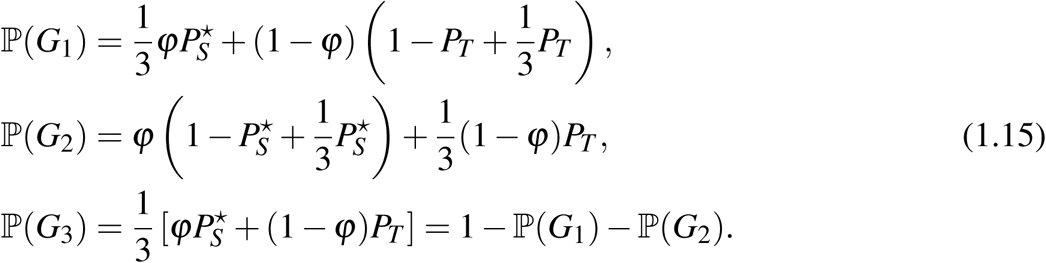

Thus ℙ(*G*_1_) < ℙ(*G*_2_), i.e., the majority-vote method infers an incorrect species tree, if and only if

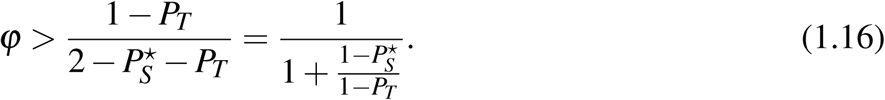

### Gene Flow under the IM Model

#### UPGMA Method with Inflow

We first consider an isolation-with-migration (IM) model with inflow. Define the migration rate (in forward-time) from species *A* to *C* under the IM model to be *M*_*AC*_ = *m*_*AC*_*N*_*C*_ migrants per generation. When we trace the genealogy of the sample backwards in time, the process of coalescence and migration during time interval (0, *τ*_*T*_) can be described by a Markov chain with three states: *S*_*abc*_, *S*_*aab*_ and *S*_*ab*_ (Hobolth *et al*., 2011; Zhu and Yang, 2012). Here *S*_*abc*_ is the initial state with the three sequences in the three populations, *S*_*aab*_ is the state after sequence *c* enters species *A* (tracing the genealogy backwards in time), with two sequences in *A* and a third in *B*, and *S*_*ab*_ is the state after sequence *c* enters *A* and coalesces with sequence *a*, so that two sequences remain in the sample.

As the model assumes unidirectional migration, the only migration possible is from *C* to *A* (when time runs backwards), at the rate *m*_*AC*_ per generation. The only coalescent possible during the time epoch (0, *τ*_*T*_) is between *a* and *c* in population *A*, at the rate 1*/*(2*N*_*A*_) per generation (Fig. 1b). Divide both rates by the mutation rate *µ* per generation so that one time unit is the expected time taken to accumulate one mutation per site. Then the migration rate becomes *w*_*AC*_ = *m*_*AC*_*/µ* = 4*M*_*AC*_*/θ*_*C*_, and the coalescent rate in species *A* becomes 2*/θ*_*A*_. Thus the backward process of migration and coalescence can be described by a Markov chain with the generator matrix *Q*

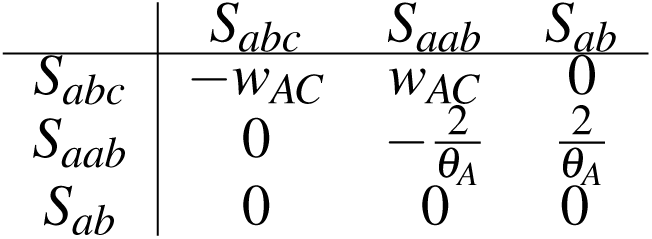

The eigenvalues of *Q* are 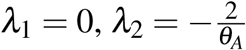, and *λ*_3_ = −*w*_*AC*_. The transition probability matrix, *P*(*t*) = exp(*Qt*), is

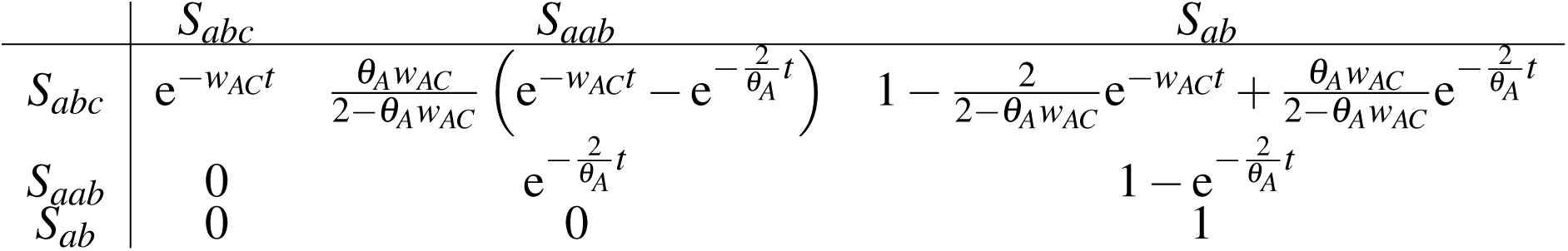

Sequences *a* and *b* can coalesce in population *R* only, with the coalescent time having an exponential distribution 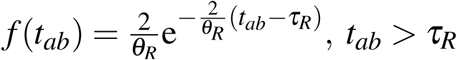, with expectation 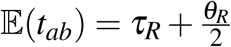. Sequences *a* and *c* can coalesce in both *A* and *R*, while sequences *b* and *c* can coalesce in both *T* and *R*. Thus 𝔼(*t*_*ab*_) > max 𝔼(*t*_*ac*_), 𝔼(*t*_*bc*_).

The probability density of coalescent time *t*_*ac*_ is

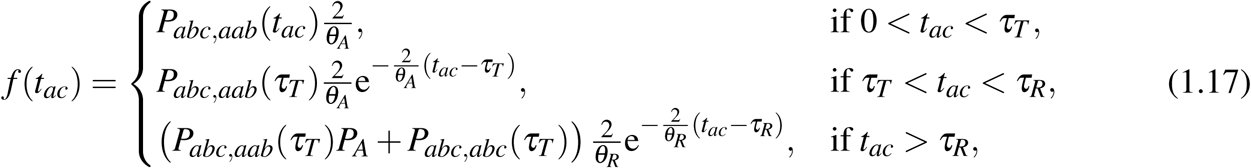

where 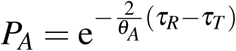 is the probability that two sequences entering population *A* at time *τ*_T_ do not coalesce before they reach time *τ*_*R*_. The expectation of *t*_*ac*_ is

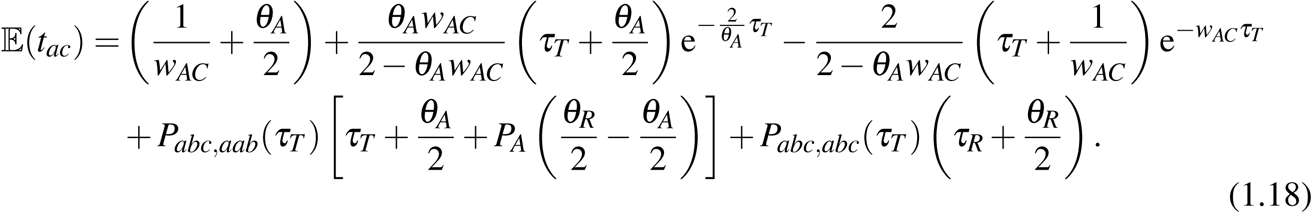

The coalescent time *t*_*bc*_ has the density

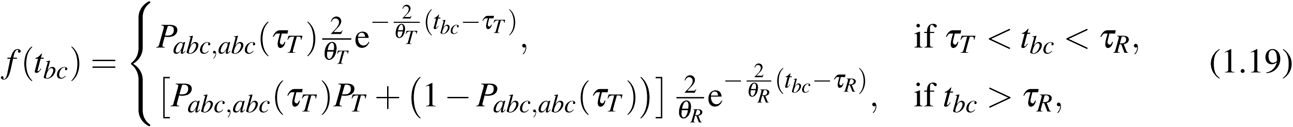

with expectation

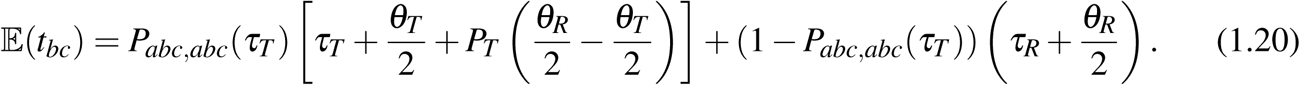

Determining the threshold value of *w*_*AC*_ or *M*_*AC*_ for which 𝔼(*t*_*bc*_) > 𝔼(*t*_*ac*_), so that the UPGMA method is inconsistent, is not analytically tractable but the threshold can be calculated numerically through a linear search for given values of parameters (*θ* s and *τ*s).

#### Majority-Vote Method with Inflow

Using the Markov chain characterization of the backward process of coalescent and migration described above, the probabilities of three gene trees can be easily derived (Fig. 1b). We have

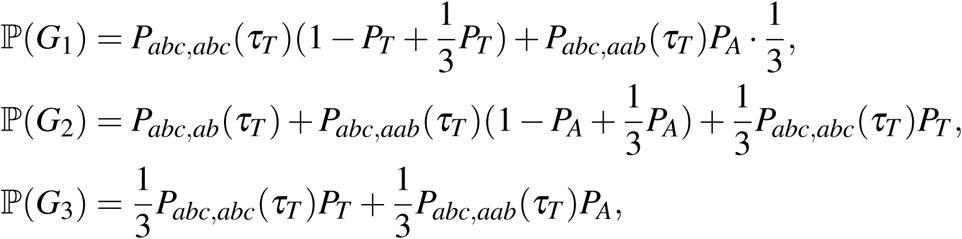

where 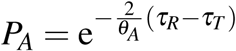. For example, gene tree *G*_1_ occurs if sequences *b* and *c* coalesce first, before two sequences coalesce. This can only occur if there is no coalescent (between sequences *a* and *c*) in the time epoch (0, *τ*_*T*_), so that the state of the Markov chain at time *τ*_*T*_ must be either *S*_*abc*_ or *S*_*aab*_. In the former case (state *S*_*abc*_ at time *τ*_*T*_), sequences *b* and *c* may coalesce first, in either species *T* (with probability 1 − *P*_*T*_) or species *R* (with probability 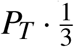). In the latter case (state *S*_*aab*_ at time *τ*_*T*_), sequences *b* and *c* may coalesce first only if there is no coalescence (between sequences *a* and *c*) in species *A* between *τ*_*T*_ and *τ*_*R*_ (with probability 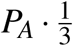).

It is easy to see that ℙ(*G*_3_) < min {ℙ(*G*_1_), ℙ(*G*_2_)*}*. We have ℙ(*G*_1_) < ℙ(*G*_2_) if and only if *Pabc,abc*(*τ*_*T*_)(2 − *P*_*T*_) + *P*_*abc,aab*_(*τ*_*T*_)*P*_*A*_ < 1, or if and only if

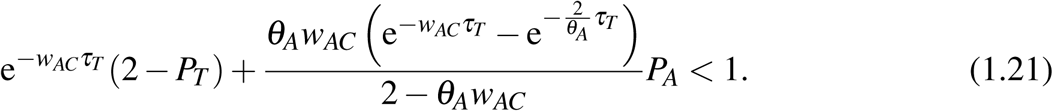

Again the threshold value of *w*_*AC*_ for this condition to be satisfied, so that the majority-vote method is inconsistent, can be calculated numerically through a linear search.

#### UPGMA Method with Outflow

Similar to the IM model of Figure 1b, we use a Markov chain to characterize the coalescent-migration process in the time interval (0, *τ*_*T*_) (Fig. 1d). The migration rate from *C* to *A* (in forward time) is *M*_*CA*_ migrants per generation or *w*_*CA*_ = *m*_*CA*_*/µ* = 4*M*_*CA*_*/θ*_*A*_ when time is scaled by the mutation rate. The coalescent rate (in population *C* after sequence *a* moves into *C*) is 1*/*(2*N*_*C*_) per generation or 2*/θ*_*C*_ on the mutational time scale. For a sample of three sequences (*a, b, c*), the three states of the Markov chain are *S*_*abc*_, *S*_*ccb*_ and *S*_*cb*_, where *S*_*ccb*_ indicates three sequences exist with two sequences in species *C* and one in *B*, and *S*_*cb*_ indicates two sequences exist with one in species *C* and one in *B*. The rate matrix *Q* is

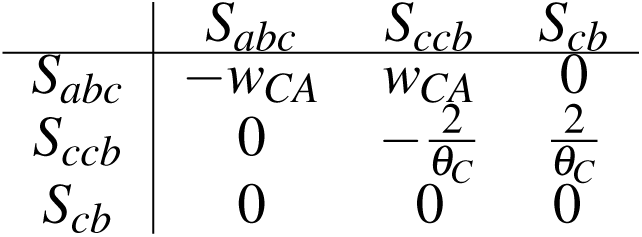

This has the eigenvalues 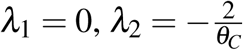, and *λ*_3_ = −*w*_*CA*_. The transition probability matrix *P*(*t*) = exp(*Qt*) is

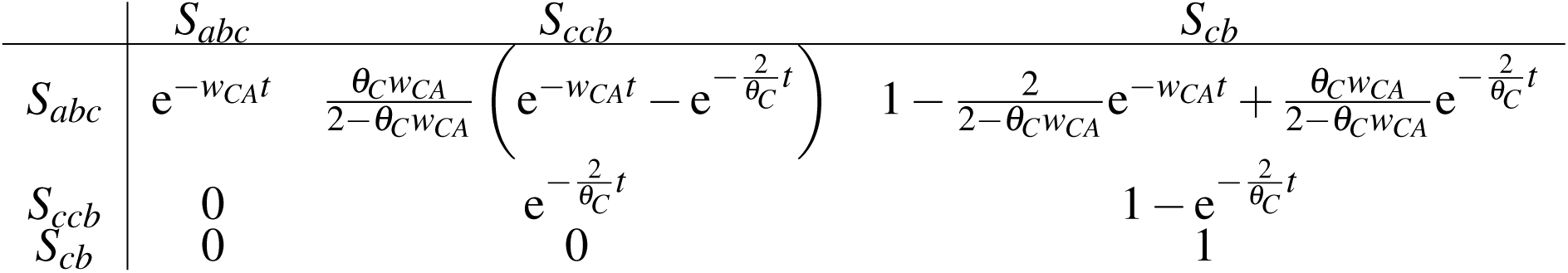

Sequences *a* and *c* can coalesce in populations *C, T* and *R*. Sequences *b* and *c* or *a* and *b* can coalesce in *T* and *R*. The probability density of coalescent time *t*_*ac*_ is

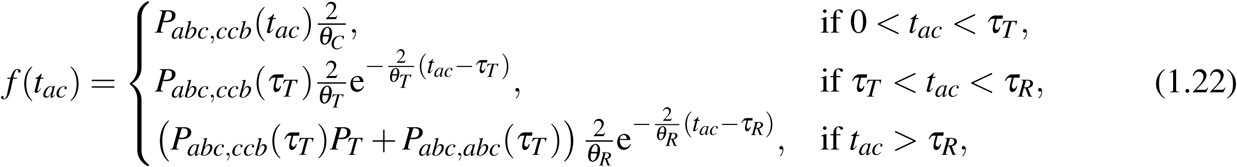

with expectation

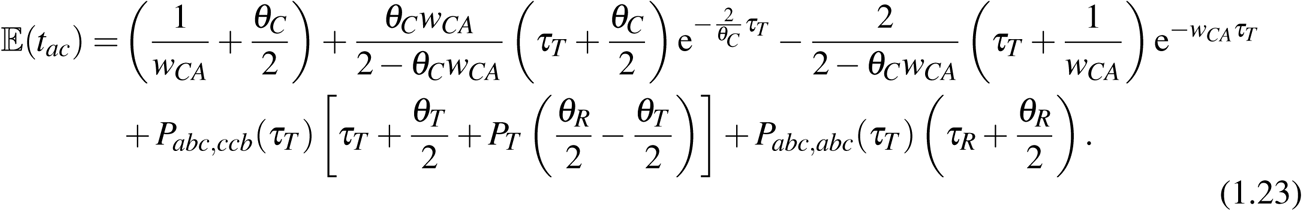

The coalescent time *t*_*bc*_ has the density

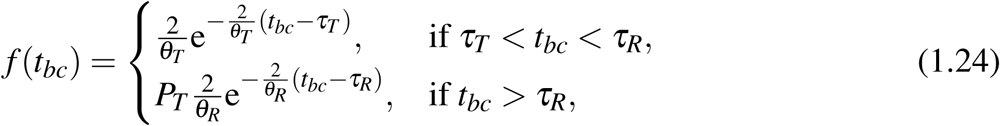

with expectation

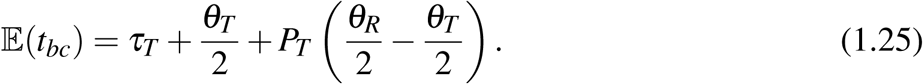

The probability density of the coalescent time *t*_*ab*_ is

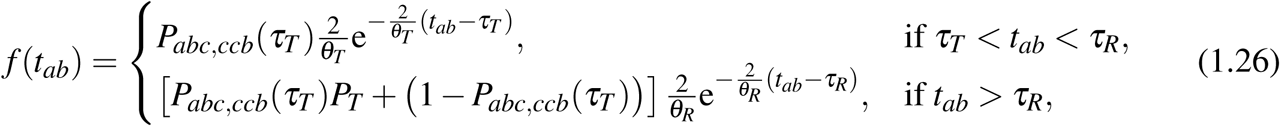

with expectation

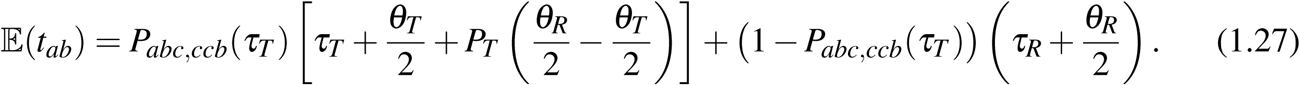

It is easy to see that 𝔼(*t*_*ab*_) > max {𝔼(*t*_*ac*_), 𝔼(*t*_*bc*_)}. Again the threshold value of *w*_*CA*_ or *M*_*CA*_ for which 𝔼(*t*_*bc*_) > 𝔼(*t*_*ac*_), so that the UPGMA method is inconsistent, can be calculated numerically through a linear search.

#### Majority-Vote Method with Outflow

The procedure we used previously to analyze the IM model with inflow is also applied here (Fig. 1d). We have

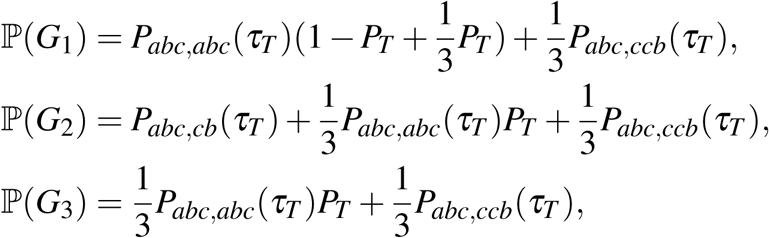

It is easy to see that *G*_3_ is the least probable gene tree, with ℙ(*G*_3_) < min {ℙ(*G*_1_), ℙ(*G*_2_)}. Furthermore, ℙ(*G*_1_) < ℙ(*G*_2_) if and only if *P*_*abc,abc*_(*τ*_*T*_)(1 − *P*_*T*_) < *P*_*abc,cb*_(*τ*_*T*_), or if and only if

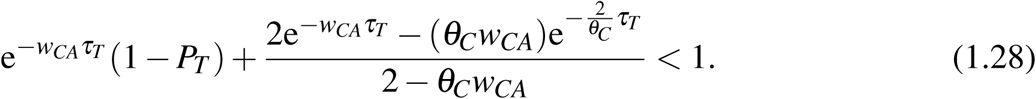

Again a linear-search algorithm can be used to determine the value of *w*_*CA*_ for which this condition is satisfied and the majority-vote method is inconsistent.

## RESULTS

### The Majority-Vote and UPGMA Methods

Here we apply the theoretical results developed above to determine the amount of gene flow, as measured by the migration rate *M* in the IM model and the introgression probability *φ* in the MSci model, that is sufficient to mislead species tree estimation. These thresholds define the boundary of the zone of inconsistency for each gene flow model and each inference method. In Figure 2, the threshold values of *φ* under the MSci model and of *M* under the IM model are plotted against *τ*_*T*_ */τ*_*R*_. All populations are assumed to have the same size (*θ*), and *τ*_*R*_ = 5*θ* or 10*θ*. In the case of the MSci model, there is an introgression event at time *τ*_*H*_ = *τ*_*S*_ = *τ*_*R*_*/*5 (see Figure 1a & c).

We focus on hard species trees with a short internal branch, that is, with *τ*_*T*_ */τ*_*R*_ much larger than 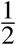 or even close to 1. Note that the plotted threshold value is the point at which the two species trees are equally good. For example, in the case of the inflow introgression model and the majority-vote method, with *τ*_*R*_ = 5*θ* and *τ*_*T*_ */τ*_*R*_ = 0.95, the threshold is *φ*_lim_ = 0.282 (Fig. 2a). Thus, if and only if the introgression probability is higher than 28.2% will the mismatching gene tree *G*_2_ be more frequent than the matching gene tree *G*_1_, and the majority-vote method infer the incorrect species tree (and be inconsistent). The threshold of *φ*_lim_ = 0.0588 for the UPGMA method (Fig. 2a) is much lower, suggesting that the UPGMA method is much more sensitive to gene flow than the majority-vote method. The results for outflow introgression are similar, with the majority-vote method being more robust to gene flow than the UPGMA method. Indeed when *τ*_*T*_ > 0.9*τ*_*R*_, the *φ* thresholds are nearly identical for inflow versus outflow introgressions.

For the inflow migration model (Fig. 2b) and with *τ*_*R*_ = 5*θ* and *τ*_*T*_ */τ*_*R*_ = 0.95, the lower limit for migration rate is *M*_*AC*_ = 0.0183 for the majority-vote method and *M*_*AC*_ = 0.00473 for UPGMA. The limiting *M*_*CA*_ values for the outflow model are similar. Again the majority-vote method is much more robust to gene flow than UPGMA. Note that in population genetic models of subdivision, gene flow of rates *M* ≈ 0.1 immigrants per generation is considered low enough so that strong population differentiation will not occur, yet such low levels can still lead to inconsistency when the species tree is difficult to reconstruct due to short internal branches.

### UPGMA Using Reconstructed Gene Trees

The gene tree probabilities we derived above are for the true gene trees. In analyses of real data gene trees estimated from sequence alignments may differ from the true gene trees due to inference errors. Here we study properties of the majority-vote method when it is applied to estimated gene trees. We expect random sampling errors to be unimportant for the UPGMA method based on average sequence distances between species because the number of sites in multi-locus datasets is huge.

The impact of phylogenetic errors under the MSC model without gene flow and in the case of three species and Jukes-Cantor (JC) model (Jukes and Cantor, 1969) was studied by Yang (2002). Without phylogenetic errors, the probabilities of the gene trees satisfy ℙ(*G*_1_) > ℙ (*G*_2_) = ℙ (*G*_3_) (Hudson, 1983). Let ℙ_*n*_(*G*_*k*_), *k* = 1, 2, 3, be the probability that the estimated gene tree (the ML gene tree, for example) is *G*_*k*_ at a locus with sequences of *n* sites, with ℙ_∞_(*G*_*k*_) = ℙ(*G*_*k*_). While phylogenetic reconstruction errors may cause the true matching gene tree (*G*_1_) to be reconstructed as a mismatching gene tree (*G*_2_ or *G*_3_) and vice versa, reconstruction errors on balance always inflate the gene tree-species tree mismatch probability, but do not change the order of the gene trees. In other words, ℙ_*n*_(*G*_1_) < ℙ (*G*_1_) and ℙ_*n*_(*G*_2_) > ℙ(*G*_2_), but the relationship ℙ_*n*_(*G*_1_) > ℙ_*n*_(*G*_2_) = ℙ_*n*_(*G*_3_) still holds (Yang, 2002). Thus the majority-vote method, when applied to estimated gene trees, is consistent, and the probability of inferring the correct species tree will approach one with the increase in the number of loci or gene trees, even if the gene trees are estimated with sampling errors. The estimate of the internal branch length (in coalescent units) in the species tree is nevertheless inconsistent, because the gene tree probabilities are distorted by phylogenetic errors.

The effects of phylogenetic errors on the gene-tree probabilities when there is gene flow (under either the IM or MSci models) are unknown. Here we use simulation to explore the issue. The ML gene tree for the case of three sequences and JC model with the molecular clock is analytically tractable (Yang, 1994, 2000). The sequence alignment at each locus can be summarized as five site-pattern counts (*n*_0_-*n*_4_), for *xxx, xxy, xyx, yxx*, and *xyz*, where *x, y, z* are any three distinct nucleotides, and the gene trees *G*_1_, *G*_2_ or *G*_3_ is the ML tree if *n*_1_, *n*_2_, or *n*_3_ is the greatest among the three. There is then no need for ML iteration to estimate the gene tree and branch lengths at each locus. We used this approach to calculate the probabilities of estimated gene trees ℙ_*n*_(*G*_1_), ℙ_*n*_(*G*_2_), and ℙ_*n*_(*G*_3_) by simulating 105 datasets or loci using BPP4 under the MSci or IM models. In Fig. 3 we choose a set of parameter values for the MSci and IM models from Fig. 2a&b for which ℙ (*G*_1_) = ℙ (*G*_2_), and simulated data at different sequence lengths. In this case phylogenetic reconstruction errors are seen to inflate the gene tree-species tree mismatch probability, with ℙ_*n*_(*G*_1_) < ℙ(*G*_1_).

We then used a linear search to find the minimum *φ* in the MSci model and minimum *M* in the IM model at which ℙ_*n*_(*G*_1_) = ℙ_*n*_(*G*_2_), with the gene tree probabilities determined by simulating 105 loci and for each locus by determining the ML tree using the observed site pattern counts. The results for the MSci model are shown in Fig. 4a&c. We focus on hard species trees and a small amount of introgression, with 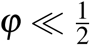. In such cases phylogenetic reconstruction errors inflate gene tree-species tree conflicts, with ℙ_*n*_(*G*_1_) < ℙ(*G*_1_). As a result, the low limit of *φ* necessary to mislead the majority-vote method of species tree estimation is lower than when true gene trees are used (Fig. 2). In other words, species tree estimation is more sensitive to introgression when gene trees are reconstructed from sequence data than when true gene trees are used. The patterns are the same for the inflow (*A* →*C*) and outflow (*C* → *A*) introgressions.

**Fig. 4.**
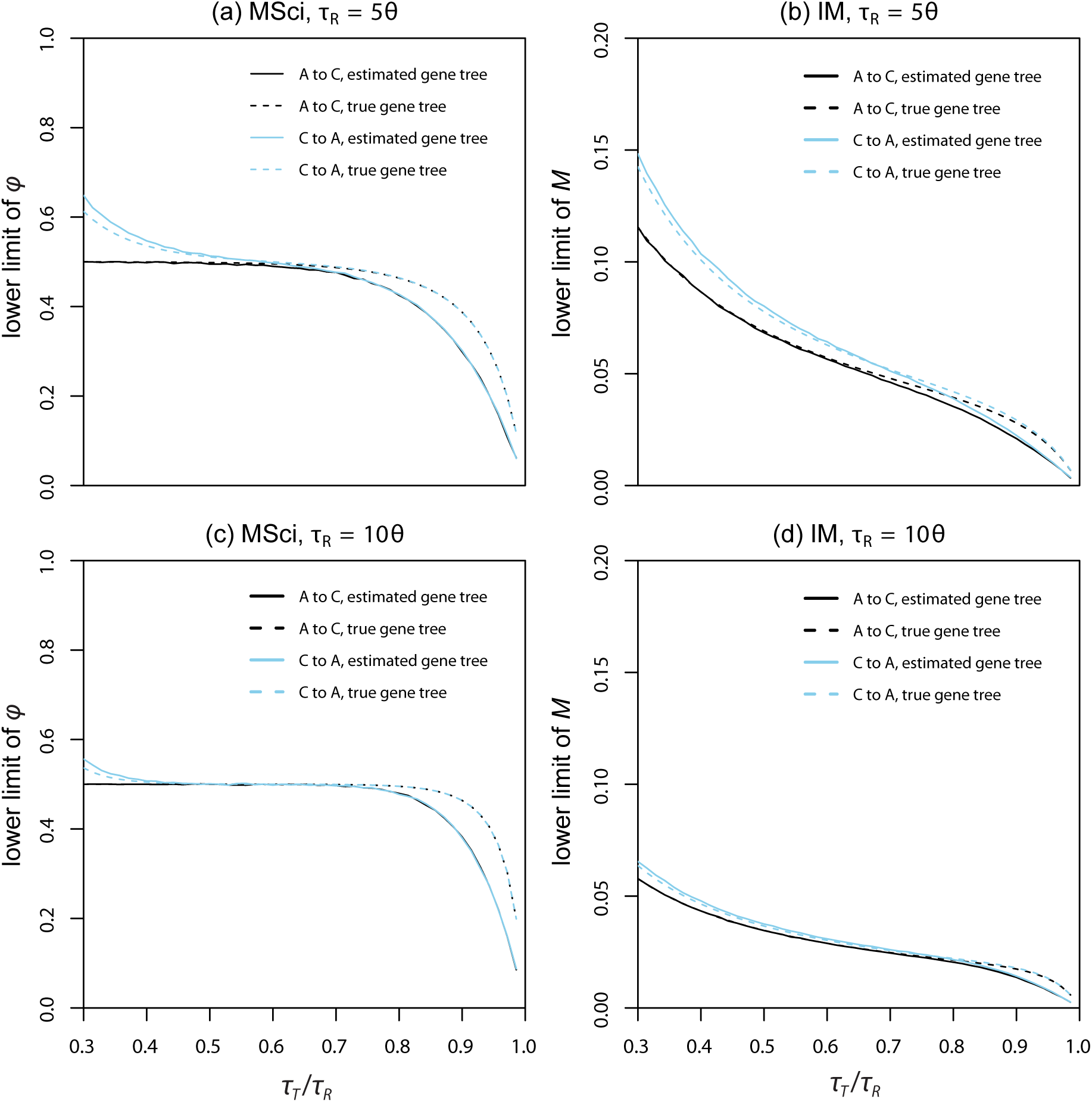
The lower limit of the introgression probability *φ* in the MSci model and the migration rate *M* in the IM model necessary to mislead the majority-vote method of species tree estimation, plotted against *τ*_*T*_ /*τ*_*R*_, when either the true or estimated gene trees are used. For estimated gene trees the sequence length is *n* = 1000. At the *φ* or *M* values shown, ℙ (*G*_1_) = ℙ (*G*_2_) when the true gene trees are used or ℙ_*n*_(*G*_1_) = ℙ_*n*_(*G*_2_) when estimated gene trees are used. The estimated gene tree is the ML tree from the sequence alignment of *n* = 1000 sites, determined by using the site pattern counts at the locus. Monte Carlo simulation, with 10^5^ replicates (loci or sequence alignments), was used to estimate the probabilities of the ML gene trees: ℙ_*n*_(*G*_1_) and ℙ_*n*_(*G*_2_). Parameters other than *φ* or *M* are fixed at *θ* = 0.01 for all populations, *τ*_*H*_ = *τ*_*S*_ = *τ*_*R*_/5 for the MSci model (a and c), while *τ*_*R*_ = 5*θ* in (a) and (b) and *τ*_*R*_ = 10*θ* in (c) and (d). Results for the true gene trees are from figure 2, shown here for comparison.

The results under the IM model are similar (Fig. 4b&d). For hard species trees with short internal branches, phylogenetic reconstruction errors inflate the gene tree-species tree conflicts, making the estimation of the species tree even harder.

### Full-Likelihood Methods

Full-likelihood methods applied to multi-locus sequence alignments, including ML and Bayesian methods, integrate over the gene tree topologies and coalescent times and naturally accommodate phylogenetic reconstruction errors due to limited number of sites at each locus (see, for a review, Xu and Yang, 2016). They are not tractable analytically. Nevertheless, in the case of three species and three sequences per locus, an efficient ML implementation of the MSC model exists in the 3S program (Yang, 2002; Dalquen *et al*., 2017). Here we use 3S to analyze datasets of 10,000 loci, assuming that at such large data size, the estimates are close to the infinite-data limits. In Figure 5, we conducted similar calculations to Figure 4, but with ML (the 3S program) replacing majority-vote. Consider 5a, where the true model is the MSci model. For each value of *τ*_*T*_, a linear search (bisection) is used to find the lowest value of *φ* in the true MSci model at which the two species trees have the same log likelihood under the MSC model, ℓ_1_ = ℓ_2_, where the log likelihood is calculated under the JC model and the molecular clock by averaging over the three gene trees and integrating over the two coalescent times in each gene tree (Yang, 2002). For each *φ* (and *τ*_*T*_ as well as other parameters in the MSci model), a dataset of 10,000 loci is simulated and analyzed using 3S to determine whether ℓ_1_ > ℓ_2_ (that is, whether *φ* is too small). Each round of bisection reduces the interval of uncertainty by a half. The scatter-points in the plots (Fig. 5) show some fluctuations, due to the finite nature of the datasets, and are used to fit a smoothed curve.

**Fig. 5.**
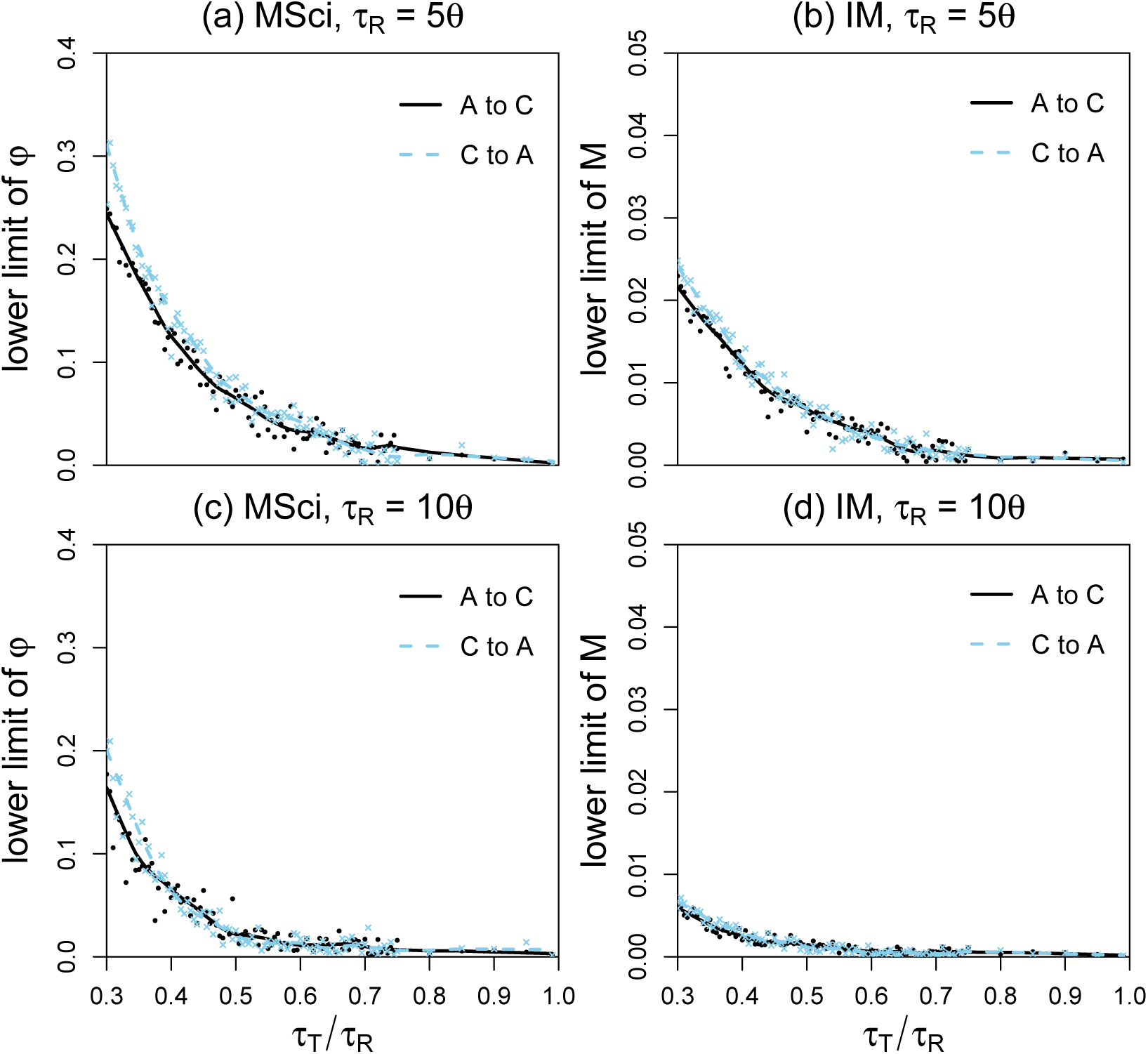
The lower limit of *φ* in the MSci model and *M* in the IM model necessary to mislead the ML/3S method of species tree estimation under the MSC, plotted against *τ*_*T*_ /*τ*_*R*_. The sequence length is 1000. Data of 10,000 loci were analyzed using 3S to determine the ML species tree. Parameter values used are the same as in Fig. 4.

The lower limits of *φ* in the MSci model and of *M* in the IM model for the ML/3S method of Figure 5 are much lower than the corresponding values for the majority-vote and the UPGMA methods of Figures 2 and 4. The ML/textsc3s method assuming MSC without gene flow infers the incorrect species tree at much lower levels of gene flow (and is less robust to gene flow) than the majority-vote or UPGMA methods.

### Differences Between the IM and MSci Models

The MSci and IM models are two idealized models that accommodate gene flow between species. The MSci model assumes episodic introgression or hybridization events that occur at fixed time points in the past, while the IM model assumes continuous-time migration with exchanges of migrants at a certain rate every generation. The MSci and IM models represent two extremes and in reality a combination of the two processes may be more realistic. When the two models are applied to the same data, a frequently asked question is how the important parameters of the two models (*φ* in MSci and *M* in IM models, say) correspond to each other. One may expect a higher migration rate *M* should correspond to a larger introgression probability *φ*, but the precise relationship may depend on the values of other parameters in the models, as well as the data configurations (the number of loci, the number of sequences, and the sequence length). Bearing in mind those caveats we suggest three analyses to address this question.

Firstly, we may compare the limiting values of *φ* under the MSci model and *M* in the IM model for the same *τ*_*T*_ */τ*_*R*_ ratio in Figure 2. For example, for the inflow migration or introgression (*A* → *C*), with *τ*_*R*_ = 5*θ* and *τ*_*T*_ */τ*_*R*_ = 0.95, the limiting values to achieve equal gene tree probabilities, P(*G*_1_) = P(*G*_2_), are *φ*_lim_ = 0.282 for the MSci model and *M*_*AC*_ = 0.0183 for the IM model. The limiting values for achieving equal average coalescent times, 𝔼(*t*_*bc*_) = 𝔼(*t*_*ac*_), are *φ*_lim_ = 0.0588 for the MSci model and *M*_*AC*_ = 0.00473 for the IM model. Here we are matching certain summaries of the data to establish a correspondence between *φ* and *M* in the two models.

Secondly, we used the IM model to simulate large datasets of 10,000 loci, with two sequences per species per locus and with sequence length of 100 or 1000 sites, and then used BPP to analyze the data under the MSci model. The rationale is that the datasets are so large that random sampling errors in the estimates are negligible, as are the impact of the prior or the differences between MLEs and Bayesian estimates. In other words, the posterior means of parameters should be close to the *pseudo-true parameter values*, which are the limits of the MLEs when the data size approaches infinity. This approach also allows us to examine the estimates of other parameters in the model besides the rate of gene flow. The results are listed in table 1.

**Table 1.** Posterior means and 95% highest probability density (HPD) credible intervals (CIs, below) for parameters in the MSci model when large datasets of 10,000 loci simulated under the IM model are analyzed using BPP under the MSci model

| $\theta_A$ | $\theta_B$ | $\theta_C$ | $\theta_R$ | $\theta_T$ | $\theta_S$ | $\theta_H$ | $\tau_R$ | $\tau_T$ | $\tau_H$ | $\varphi$ |
| --- | --- | --- | --- | --- | --- | --- | --- | --- | --- | --- |
| Inflow migration ($A \rightarrow C$) with $M_{AC} = 0.0393$, $n = 100$ sites | | | | | | | | | | |
| 0.0096 | 0.0095 | 0.0111 | 0.0121 | 0.0185 | 0.0149 | 0.0106 | 0.0478 | 0.0386 | 0.0075 | 0.3319 |
| 0.0093 | 0.0093 | 0.0107 | 0.0109 | 0.0131 | 0.0140 | 0.0092 | 0.0471 | 0.0372 | 0.0072 | 0.3208 |
| 0.0099 | 0.0098 | 0.0115 | 0.0134 | 0.0241 | 0.0157 | 0.0119 | 0.0485 | 0.0401 | 0.0077 | 0.3435 |
| Inflow migration ($A \rightarrow C$) with $M_{AC} = 0.0393$, $n = 1000$ sites | | | | | | | | | | |
| 0.0091 | 0.0099 | 0.0100 | 0.0106 | 0.0115 | 0.0250 | 0.0100 | 0.0496 | 0.0398 | 0.0050 | 0.4393 |
| 0.0089 | 0.0097 | 0.0098 | 0.0102 | 0.0107 | 0.0243 | 0.0095 | 0.0495 | 0.0395 | 0.0049 | 0.4300 |
| 0.0094 | 0.0101 | 0.0103 | 0.0109 | 0.0122 | 0.0256 | 0.0106 | 0.0498 | 0.0400 | 0.0051 | 0.4485 |
| Outflow migration ($C \rightarrow A$) with $M_{CA} = 0.0419$, $n = 100$ sites | | | | | | | | | | |
| 0.0108 | 0.0098 | 0.0096 | 0.0154 | 0.0032 | 0.0122 | 0.0094 | 0.0437 | 0.0417 | 0.0068 | 0.3012 |
| 0.0104 | 0.0095 | 0.0092 | 0.0145 | 0.0020 | 0.0115 | 0.0083 | 0.0431 | 0.0410 | 0.0066 | 0.2907 |
| 0.0112 | 0.0101 | 0.0099 | 0.0163 | 0.0049 | 0.0129 | 0.0106 | 0.0442 | 0.0424 | 0.0071 | 0.3122 |
| Outflow migration ($C \rightarrow A$) with $M_{CA} = 0.0419$, $n = 1000$ sites | | | | | | | | | | |
| 0.0102 | 0.0099 | 0.0094 | 0.0111 | 0.0094 | 0.0243 | 0.0098 | 0.0485 | 0.0401 | 0.0051 | 0.4437 |
| 0.0099 | 0.0097 | 0.0091 | 0.0107 | 0.0089 | 0.0237 | 0.0093 | 0.0483 | 0.0399 | 0.0050 | 0.4342 |
| 0.0105 | 0.0101 | 0.0096 | 0.0115 | 0.0099 | 0.0250 | 0.0104 | 0.0488 | 0.0403 | 0.0052 | 0.4533 |

Note.— The true model is IM, with parameter values $\theta_A = \theta_B = \theta_C = \theta_T = \theta_R = 0.01$ , $\tau_R = 0.05$ and $\tau_T = 0.04$ , and with $M_{AC} = 0.0393$ and $M_{CA} = 0.0419$ . These are the lower limits of $M$ in the IM model, at which $\mathbb{P}(G_1) = \mathbb{P}(G_2)$ (Fig. 2). Large datasets of 10,000 loci, with two sequences from each species per locus and with the sequence length $n = 100$ or 1000 sites, were simulated under the IM model, and analyzed using BPP under the MSci model. Inverse-gamma priors are assigned on $\theta$ and $\tau_0$ : $\theta \sim \text{IG}(3, 0.02)$ with mean $0.02/(3-1) = 0.01$ for all $\theta$ s, and $\tau_0 \sim \text{IG}(3, 0.1)$ with mean $0.1/(3-1) = 0.05$ for the age of the species-tree root, while the introgression probability $\varphi$ is assigned the $\mathbb{U}(0, 1)$ prior.

In the case of inflow migration (*A* → *C*), population sizes (*θ*) for most populations (in particular, the extant species) are well estimated, especially at *n* = 1000. Species divergence times *τ*_*R*_ and *τ*_*T*_ estimated under MSci are very similar to the true values under the IM model (0.05 and 0.04, respectively). The age of the hybridization node *τ*_*H*_ (= *τ*_*S*_) is very small, especially at the larger sequence length. Under the IM model, migration occurs over the whole time interval (0, *τ*_*T*_), and one might naively expect *τ*_*H*_ under MSci to be close to the mid value. Instead the MSci estimate of *τ*_*H*_ is near the lower limit. Indeed the estimate should be smaller for longer sequences and/or more loci, because under the MSci model, the hybridization time must be smaller than the between-species sequence divergence time, with *τ*_*H*_ < *t*_*ac*_. Thus, longer sequences or more loci should provide stronger evidence that the minimum sequence divergence *t*_*ac*_ generated under the IM model can be arbitrarily small, leading to reduced estimates of *τ*_*H*_ under the MSci model. The migration rate *M*_*AC*_ = 0.0393 in the IM model is relatively small, while the estimates of *φ* are substantial, at 33-44%. The results for the case of outflow migration (*C→ A*) are similar to those of inflow migration (table 1). Again, species divergence times *τ*_*R*_ and *τ*_*T*_ are well estimated, as are population sizes, even though the model is incorrect. The estimated hybridization time *τ*_*H*_ (= *τ*_*S*_) is very small. Although the migration rate is only *M*_*AC*_ = 0.0419, the estimates of *φ* in the MSci model are substantial, at 30-44%. Overall, very low migration rates, on the order of *M* = 1-5% may correspond to substantial introgression probabilities with *φ* in the range of 20-50%.

Thirdly, we analyzed the case of two species (*A* and *B*) with one sequence per locus for each species when the true sequence distance (or the coalescent time between the two sequences) is known (Fig. 6). Using the theory for the IM model developed earlier, we have the probability density of coalescent time *t* between the two sequences to be

**Fig. 6.**
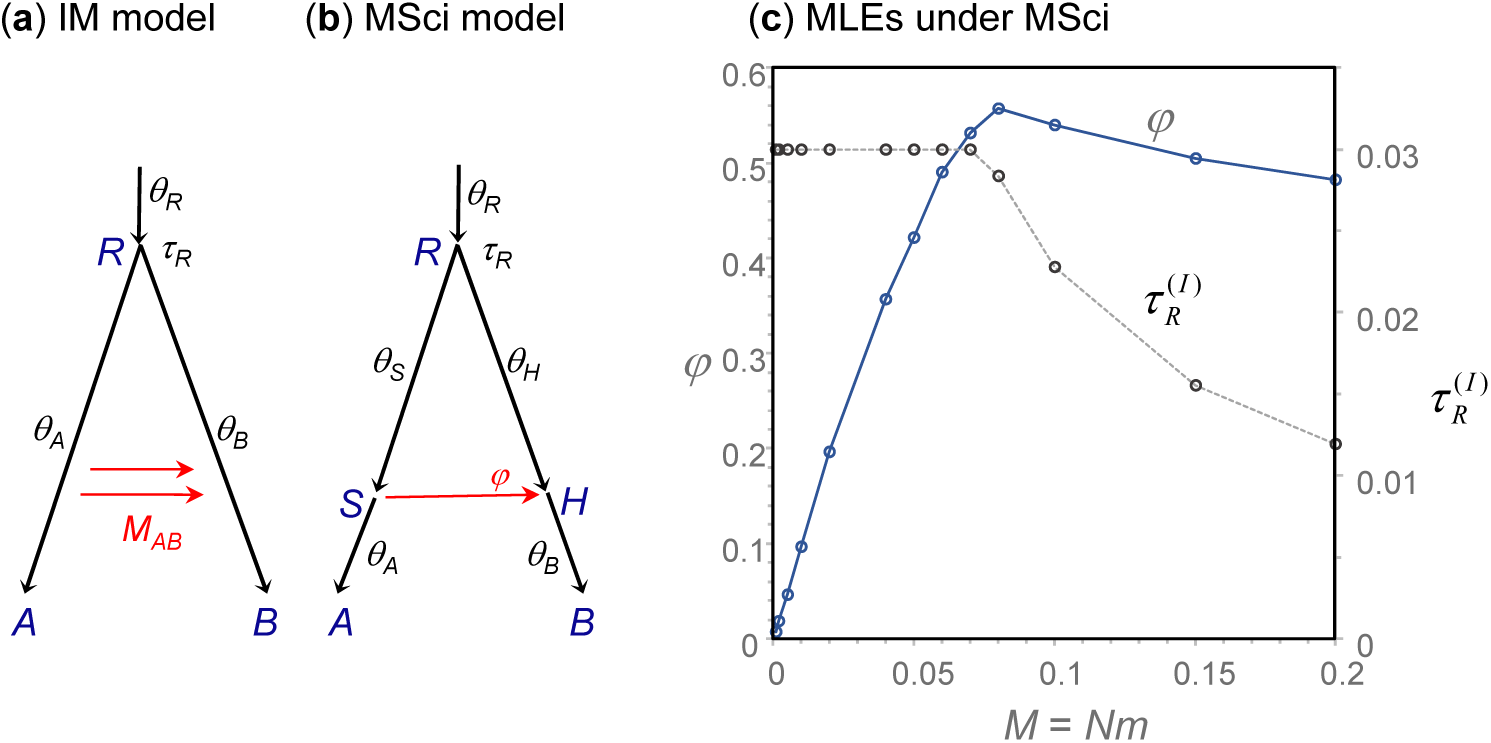
The (**a**) migration and (**b**) introgression models for two species (*A, B*), and (**c**) the pseudo-true parameter values of *φ* and 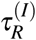 in the introgression model plotted against the migration rate *M*_*AB*_(= *wθ_B_/*4). The pseudo-true parameter values are the limiting values of the MLEs under the MSci model when the amount of data (the number of loci) approaches infinity. The true model is the IM model and the data (the distribution of the coalescent time *t*) is fitted using the MSci model. Other parameters in the migration model are fixed at 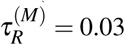 and *θ* = 0.01 (for all populations). In the MSci model, *θ* = 0.01 is assumed for all populations, while *φ* and 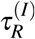 are estimated by minimizing the KL divergence (equation 1.31).

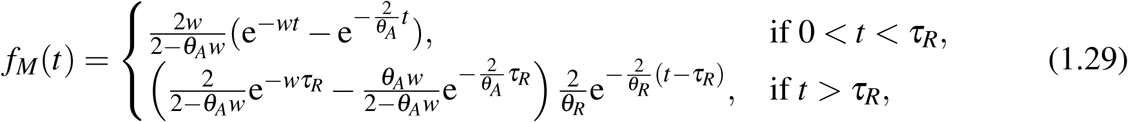

where *w* = *m*_*AB*_*/µ* = 4*M*_*AB*_*/θ*_*B*_ is the mutation-scaled migration rate from *A* to *B*. The density depends on *w* but not on *M*_*AB*_ and *θ*_*B*_ individually, so that the parameters specifying the density for the migration model are ***θ***^(*M*)^ = *{w, θ*_*A*_, *θ*_*R*_, *τ*_*R*_*}*. Similarly the density under the introgression model is

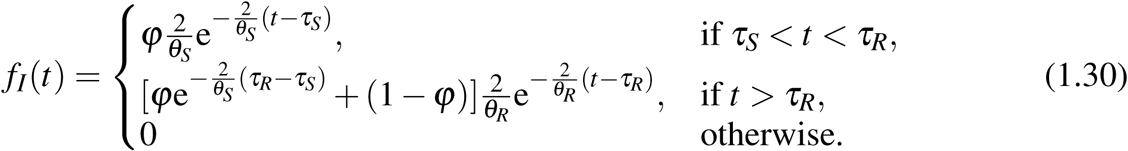

This is specified by parameters ***θ***^(*I*)^ = {*ϕ, θ*_*S*_, *θ*_*R*_, *τ*_*H*_, *τ*_*R*_}.

The Kullback-Leibler (KL) divergence

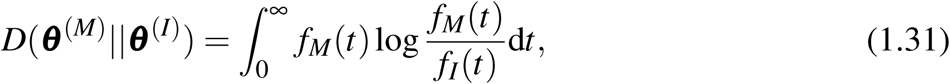

is a measure of distance from the fitting introgression model to the true migration model. By minimizing *D*, we obtain the pseudo-true parameter values under MSci: the limiting values of the MLEs when the amount of data (the number of loci) approaches infinity, when the true model is the IM model with parameters ***θ***^(*M*)^. Here ***θ***^(*M*)^ are fixed while ***θ***^(*I*)^ are being optimized. Because the IM model allows arbitrarily small coalescent time *t* while *τ*_*H*_ > *t* under the MSci model, we have *τ*_*H*_ = 0. Note that some parameters (such as *τ*_*R*_ and *θ*_*R*_) are common between the two models but they may take different values: when the parameter definition may be unclear from the context, we use 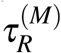, with superscripts ‘(*M*)’ or ‘(*I*)’, to indicate the model involved.

Figure 6c shows the estimates of *φ* and 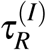 under the assumption that *θ* is the same for all populations and also between the two models; thus *θ* = 0.01 is fixed and only *φ* and 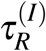 are estimated by minimizing *D* in equation 1.31. We applied Gaussian quadrature to calculate the integral of equation (1.31), dealing with the discontinuity in the integrand (see Fig. S1 for details). When *M* in the IM model is small, *φ* increases nearly linearly with the increase of *M* (Fig. 6c). However, when *M* is large (> 0.08, say, corresponding to *φ* = 0.56) and increases further, *φ* decreases. We note that when *φ* > 0.5, the biological interpretations of the model or parameter change.

As a summary of all three analyses above, we note that small values of the migration rate, in the order of 0.01-0.1 migrants per generation, may correspond to large introgression probabilities. Our analyses also suggest that the IM and MSci models make very different predictions of the distribution of coalescent times, so that the two models may be easily distinguishable using genomic sequence data when both models are implemented in the same program.

### African Mosquito Data Example

The *Anopheles gambiae* species complex is comprised of eight recognized species and includes major malaria vectors in Africa. Genome sequence data from six of the species, *A. gambiae* (G), *A. coluzzii* (C), *A. arabiensis* (A), *A. melas* (L), *A. merus* (R), and *A. quadriannulatus* (Q), have been analyzed to estimate the species phylogeny and to infer the direction and intensity of gene flow across species (Fontaine *et al*., 2015; Thawornwattana *et al*., 2018). *A. gambiae, A. coluzzii*, and *A. arabiensis* have large overlapping geographical distributions across sub-Saharan Africa and are major malaria vectors (Wiebe *et al*., 2017). *A. melas* and *A. merus* are found in coastal waters of eastern and western Africa, respectively, and are minor vectors. *A. quadriannulatus* does not bite humans and is not a malaria vector.

Fontaine *et al*. (2015) analyzed the genomic sequence data and suggested that gene flow is so widespread that the predominant gene trees for the autosomes are different from the species phylogeny, and that the X chromosome, which is not affected by gene flow, reflects the true species history. This conclusion is supported in the re-analysis of Thawornwattana *et al*. (2018), although the inferred species trees were different. Here we use the species tree of Thawornwattana *et al*. (2018) (Fig. 7) and the parameter estimates from Flouris *et al*. (2019) to confirm that under the MSci model the introgression rates affecting the autosomes are high enough to mislead species tree estimation.

**Fig. 7.**
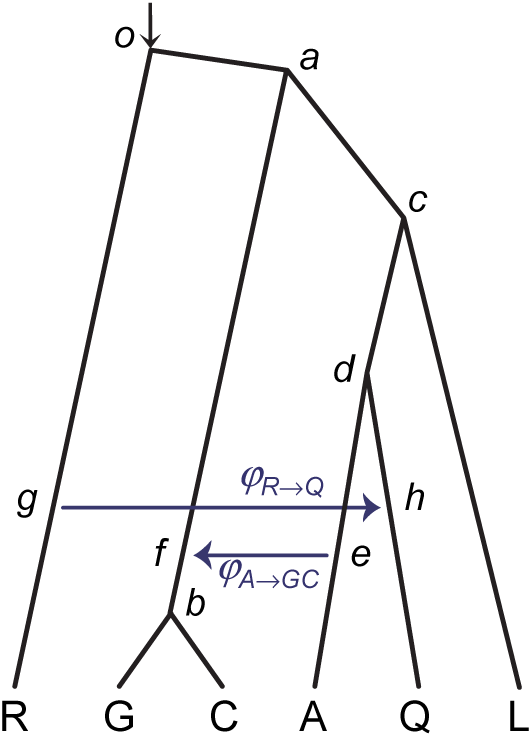
Species tree with two migration events for the *Anopheles gambiae* species complex. Redrawn following (Thawornwattana *et al*., 2018, Fig. 6).

For the A → GC introgression, we used parameter estimates (*τ*s and *θ* s) for the triplet AQG, to determine the low limit of *φ* in the MSci model. The results are shown in table 2. For both the coding and noncoding data and for almost all chromosomal arms, estimates of *φ* are larger than the limiting value *φ*_lim_. Thus, both majority-vote (based on gene tree topologies) and UPGMA (based on average sequence divergences) are inconsistent and are expected to infer an incorrect species tree.

**Table 2.**
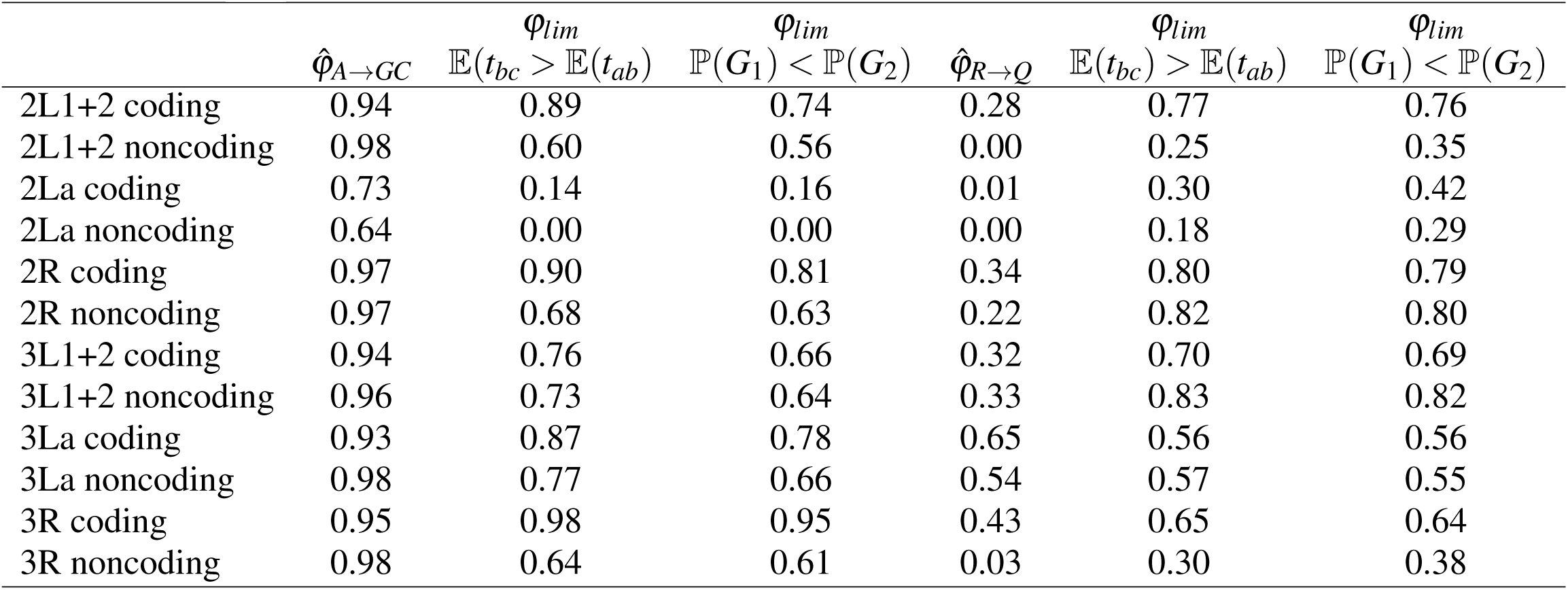
Estimates and limiting values of *φ* for different chromosomal arms obtained from the genomic data of the *Anopheles gambiae* species complex

| | $\hat{\phi}_{A \rightarrow GC}$ | $\phi_{lim}$<br>$\mathbb{E}(t_{bc}) > \mathbb{E}(t_{ab})$ | $\phi_{lim}$<br>$\mathbb{P}(G_1) < \mathbb{P}(G_2)$ | $\hat{\phi}_{R \rightarrow Q}$ | $\phi_{lim}$<br>$\mathbb{E}(t_{bc}) > \mathbb{E}(t_{ab})$ | $\phi_{lim}$<br>$\mathbb{P}(G_1) < \mathbb{P}(G_2)$ |
| --- | --- | --- | --- | --- | --- | --- |
| 2L1+2 coding | 0.94 | 0.89 | 0.74 | 0.28 | 0.77 | 0.76 |
| 2L1+2 noncoding | 0.98 | 0.60 | 0.56 | 0.00 | 0.25 | 0.35 |
| 2La coding | 0.73 | 0.14 | 0.16 | 0.01 | 0.30 | 0.42 |
| 2La noncoding | 0.64 | 0.00 | 0.00 | 0.00 | 0.18 | 0.29 |
| 2R coding | 0.97 | 0.90 | 0.81 | 0.34 | 0.80 | 0.79 |
| 2R noncoding | 0.97 | 0.68 | 0.63 | 0.22 | 0.82 | 0.80 |
| 3L1+2 coding | 0.94 | 0.76 | 0.66 | 0.32 | 0.70 | 0.69 |
| 3L1+2 noncoding | 0.96 | 0.73 | 0.64 | 0.33 | 0.83 | 0.82 |
| 3La coding | 0.93 | 0.87 | 0.78 | 0.65 | 0.56 | 0.56 |
| 3La noncoding | 0.98 | 0.77 | 0.66 | 0.54 | 0.57 | 0.55 |
| 3R coding | 0.95 | 0.98 | 0.95 | 0.43 | 0.65 | 0.64 |
| 3R noncoding | 0.98 | 0.64 | 0.61 | 0.03 | 0.30 | 0.38 |

For the R → Q introgression, we use the RQA triplet. All estimates of *φ*_*R*→*Q*_ are smaller than *φ*lim for both the majority-vote and UPGMA methods except for the 3La coding region. Thus, for most of the autosomal arms introgression from R to Q did not reach a sufficient intensity to mislead species tree estimation.

## DISCUSSION

### The Impact of Gene Flow on Species Tree Estimation

The impact of gene flow, either in the form of episodic introgressive hybridisation or continuous migration, on species tree estimation clearly depends on how challenging the species tree is. Our analyses suggest that when the species tree is hard with very short internal branch lengths, even a small amount of gene flow can cause species tree estimation to become inconsistent. We found that the limiting *φ* and *M* values for the gene tree probabilities are much higher than the values for the sequence distances (Figures 2 and 4), indicating that the majority vote method based on gene tree topologies is more robust to gene flow than the UPGMA method based on average sequence distances. The full likelihood method making use of information in both gene tree topologies and branch lengths is even more sensitive. This difference in sensitivity may be explained by the fact that a small amount of gene flow may easily affect the branch lengths or sequence distances but may not alter the distribution of gene tree topologies. For hard species trees, phylogenetic reconstruction errors tend to inflate the gene tree-species tree conflicts, adding further challenges to correct inference of the species tree.

When the effects of gene flow is a concern, it is important to use species tree methods that account for both the coalescent process and cross-species gene low. Such methods are under active development, including methods for testing gene flow using summaries of sequence data (Green *et al*., 2010; Durand *et al*., 2011; Yu and Nakhleh, 2015) and full-likelihood methods based on sequence alignments (Hey and Nielsen, 2004; Dalquen *et al*., 2017; Hey *et al*., 2018; Wen and Nakhleh, 2018; Zhang *et al*., 2018). Furthermore, we have in this paper examined the effects of gene flow on the inference of species tree topology only. The impact of gene flow on evolutionary parameters such as species divergence times merits detailed study (Wen and Nakhleh, 2018).

### The Relationships between Anomaly Zone and Consistency Zone

The set of species tree and parameter values under the MSC model for which the most probable gene tree has a topology that differs from the species tree is called the anomaly zone (Degnan and Rosenberg, 2006). Some authors appear to believe that all species tree methods are inconsistent in the anomaly zone. For example, Degnan and Rosenberg (2006) wrote that “the use of multiple genomic regions for species tree inference is subject to a surprising new difficulty, the problem of ‘anomalous gene trees’.” We note that full likelihood methods based on sequence alignments, including both ML and Bayesian inference, are consistent over the whole parameter space, both inside and outside the anomaly zone (Xu and Yang, 2016). Degnan and Rosenberg (2006) made suggestions to overcome the obstacle of anomalous gene trees, including sampling multiple individuals from the same species and analyzing and assembling species triplets (as anomalous gene trees do not exist in the case of three species); it should be mentioned that standard statistical methods based on the likelihood function do not suffer from the obstacle and should indeed be the method of choice.

Similarly Solis-Lemus *et al*. (2016) explored the anomaly zone in the presence of gene flow and concluded that “we identified a situation where the presence of gene flow causes the appearance of AUGTs [anomalous unrooted gene trees] on four taxa, and anomalous rooted gene trees on three taxa. In this situation with only four (or three) taxa, all species tree methods are necessarily inconsistent: a coalescent model ignoring gene flow would necessarily favor one of the two incorrect four-taxon unrooted trees (or three-taxon rooted trees), when the true species tree is the one that is supported by the least proportion of genes.” The authors’ conclusion that “all species tree methods are necessarily inconsistent” in the anomaly zone is not correct. Different species tree methods have different statistical properties and the anomaly zone is the zone of inconsistency for the majority-vote method (using true trees) only; other methods have different zones of inconsistency.

In the situations studied in this paper there is gene flow but the inference methods ignore it so that the model is violated. In this case, full likelihood methods may be inconsistent, just like other methods. For both the inflow (*A* →*C*) and outflow (*C* → *A*) models of gene flow, and for both the IM and MSci models, the majority-vote method based on gene tree topologies has a smaller zone of inconsistency than the UPGMA method based on average sequence distances (Fig. 2), and full likelihood methods ignoring gene flow have a larger zone of inconsistency than either the UPGMA or majority-vote methods.

## 2. ACKNOWLEDGEMENTS

This study has been supported by Biotechnological and Biological Sciences Research Council grants (BB/N000609/1 and BB/P006493/1) to Z.Y.

---

**Figure S1.**
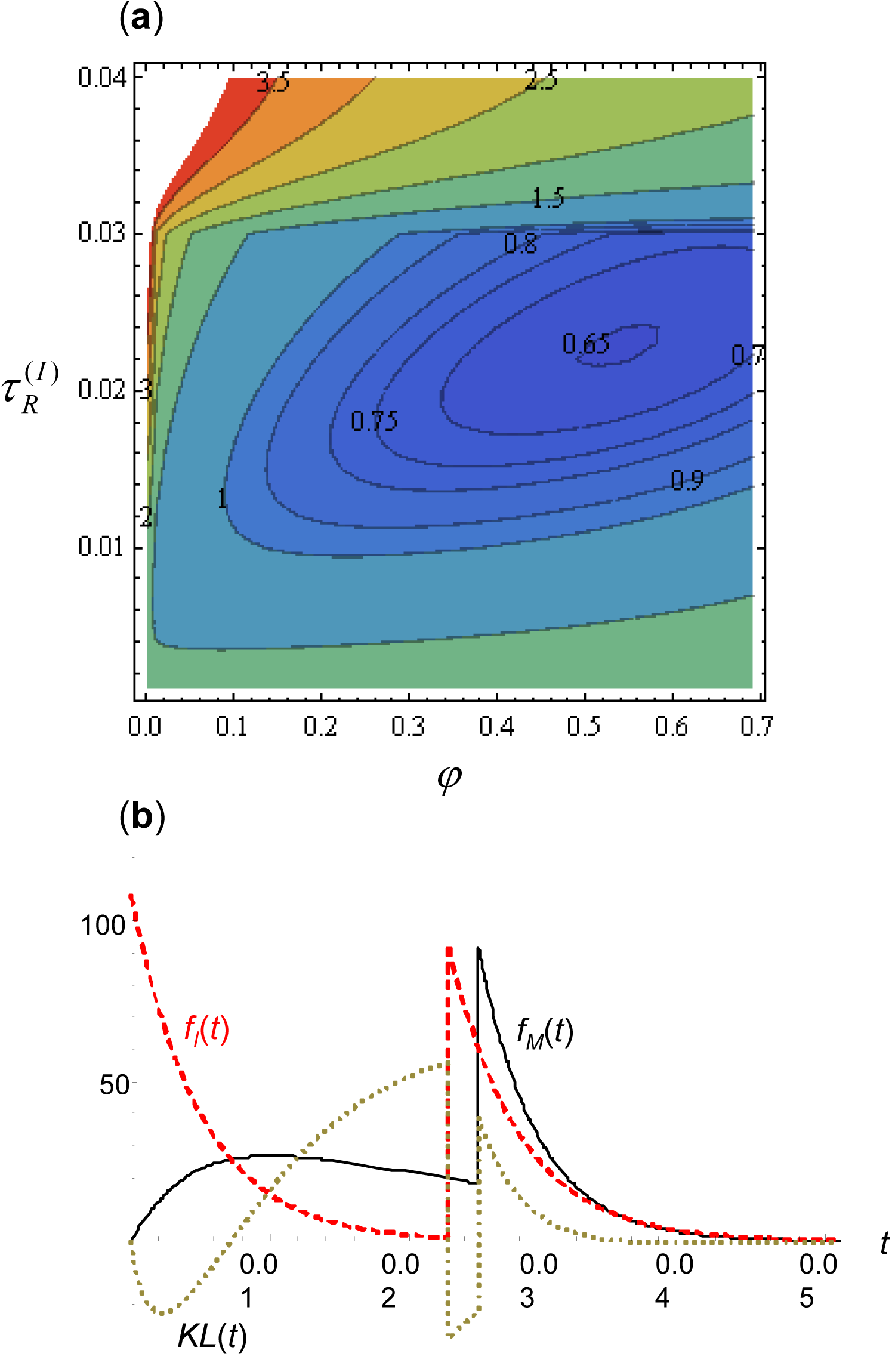
(**a**) Contour plot of *D*(*θ*_M_ ‖ *θ*_I_) as a function of *φ* and *τ*_*R*_ in the introgression model when the true model is the migration model with *M* = 0.1 (or *w* = 40). Other parameters in the migration model are *τ*_*R*_ = 0.03 for the age of the root and *θ* = 0.01 for all populations. The MLE under the Msci model, which minimizes *D*, is at *φ* = 0.5395 and *τ*_*R*_ = 0.02280. Note that the surface is not smooth at *τ*_*R*_ = 0.03. (**b**) The density of coalescent time *fM*(*t*) under the IM model and *fI*(*t*) under the Msci model as well as the integrand, *fM*(*t*) log *fM*(*t*)/*fI*(*t*), of equation 31. The parameters in the IM model are *M* = 0.1 (or *w* = 40) and *τ*_*R*_ = 0.03, and *θ* = 0.01 for all populations. The parameters under the Msci model are *φ* = 0.5395 and *τ*_*R*_ = 0.022803, while *θ* = 0.01 is assumed for all populations: these *φ* and *τ*_*R*_ values are the best-fitting values under the MSci model when the true model is the IM model. Note that the contour plot in (**a**) is not smooth at 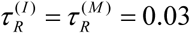, and the integrand in (**b**) has two discontinuity points at 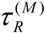 and 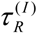.

